# Blackbird: structural variant detection using synthetic and low-coverage long-reads

**DOI:** 10.1101/2024.11.17.624011

**Authors:** Dmitry Meleshko, Rui Yang, Salil Maharjan, David C. Danko, Anton Korobeynikov, Iman Hajirasouliha

## Abstract

**Motivation:** Recent benchmarks of structural variant (SV) detection tools revealed that the majority of human genome structural variations (SVs), especially the medium-range (50-10,000 bp) SVs cannot be resolved with short-read sequencing, but long-read SV callers achieve great results on the same datasets. While improvements have been made, high-coverage long-read sequencing is associated with higher costs and input DNA requirements. To decrease the cost one can lower the sequence coverage, but the current long-read SV callers perform poorly with coverage below 10×. Synthetic long-read (SLR) technologies hold great potential for structural variant (SV) detection, although utilizing their long-range information for events smaller than 50 kbp has been challenging.

**Results:** In this work, we propose a hybrid novel integrated alignment- and local-assembly-based algorithm, Blackbird, that uses SLR together with low-coverage long reads to improve SV detection and assembly. Without the need for a computationally expensive whole genome assembly, Blackbird uses a sliding window approach and barcode information encoded in SLR to accurately assemble small segments and use long reads for an improved gap closing and contig assembly. We evaluated Blackbird on simulated and real human genome datasets. Using the HG002 GIAB benchmark set, we demonstrated that in hybrid mode, Blackbird demonstrated results comparable to state-of-the-art long-read tools, while using less long-read coverage. Blackbird requires only 5× coverage to achieve F1 scores (0.835 and 0.808 for deletions and insertions) similar to PBSV (0.856 and 0.812) and Sniffles2 (0.839 and 0.804) using 10× Pacbio Hi-Fi long-read coverage.

## 1 Introduction

Structural variations (SVs) greatly contribute to human genome diversity [12]. Characterizing the full spectrum of SVs in human genomes makes it possible to understand the distribution of SVs in different populations [8]. Additionally, SVs have been found to play a critical role in the development of certain diseases, such as cancer [22] and Alzheimer’s [36]. SVs are typically referred to any genome sequence altering event, except for deletions or insertions (indels) shorter than 50 bp [32]. Structural variants (SVs) are often classified into two categories: (a) large-scale events affecting tens of thousands of bases in the genome, such as translocations, large inversions, and segmental duplications, and (b) smaller events, less than 10 kbp in size, which can be classified as insertions, deletions, or other types. Notable examples of the latter class include mobile element insertions and deletions. Both classes of SVs are extensively studied using various sequencing technologies and algorithms. Currently the vast majority of whole genome sequencing is performed using standard short-read technologies. While SVs are biologically and clinically important, their identification remains challenging. Short-read methods have low sensitivity and can introduce false-positive calls due to their relatively short fragment lengths and lack of long-range information. [6, 33]. In particular, prediction of SV events inside or around repetitive regions using short reads is very challenging. All available SV calling tools are based on the assumption that read mapping to the reference genome is correct, however, in a typical sequencing experiment, less than 80% of the reads are mapped unambiguously to the reference genome. For ambiguous reads originating from repeat sequences, often only one random or best hit is considered, which makes accurate SV prediction extremely challenging [33]. Due to the low cost of short-read sequencing, however, multiple tools such as SVaBa [35], LUMPY [20], Manta [7], GRIDSS [4], Tardis [15], Cue [27] and others are developed and being used for genome-wide SV calling. These methods use different signatures of short-read alignments to the reference genome in order to characterize SV events. For example, DELLY uses read-pair and split-read signatures to identify SV breakpoints. LUMPY utilizes split-read, read pair, and prior knowledge about SVs (e.g. a database of common SVs) and combines these signals into a probabilistic framework to calculate the probability of novel break-point for a given position. SVaBa uses a local assembly approach, that allows calling insertions with higher sensitivity. Cue uses a deep learning-based approach to call and genotype SVs [27]. Cue encodes various SV signals from read alignments into 2D images and uses a neural network for SV detection.

Recent technology advancements, specially increases in accuracy and decreases in cost of long-read platforms (PacBio, Oxford Nanopore) promise to overcome some of the limitations of short-reads. Long-read methods such as SMRT-SV [5, 16], NextSV [11], Sniffles [30] or SVIM [14] can effectively identify most SV types, including long insertions, by aligning long reads to the reference genome and subsequently sub-assembling regions with breakpoint signatures. However, high-coverage long-read sequencing may still be more expensive compared to short-read methods. Additionally, long reads require high-quality and high-input DNA. One way to offset the cost of SV calling using long-read sequencing is to lower the sequence coverage. However, existing tools typically experience a rapid decline in recall rates when coverage falls below a certain threshold.

Recently introduced low-cost and low-input synthetic long-read (SLR) sequencing technologies—such as single tube long fragment read (stLFR) and UST’s Transposase Enzyme-Linked Long-read Sequencing (TELL-Seq) provide long-range information useful for characterizing genomes and metagenomes [38]. In these new technologies, input DNA is first sheared into long fragments of 5–100 kbp. After shearing, a barcode is ligated to short reads from these fragments, ensuring that short reads originating from the same fragment share the same barcode. Finally, the short reads are sequenced using standard sequencing technologies (e.g., Illumina HiSeq).

Despite their potential, synthetic long-read (SLR) methods have not gained widespread popularity. This is partly due to a lack of sensitive, robust, and scalable methods that leverage SLR technology. To date, only a few studies have attempted to compare SLR with conventional shortread or long-read methods for structural variant (SV) detection [25, 27].

Existing SLR-specific SV detection methods such as NAIBR [9], Linked-SV [11], or GROC-SVs [31] utilize barcode similarity between distant parts of the target genome, therefore they are able to call only large events. Given that SLR platforms provide only shallow coverage of long fragments per barcode, it is challenging to detect smaller events using barcode-specific information, since it is hard to distinguish between intramolecule gap between consecutive short reads and real SV event. An alternative approach for genome-wide SV detection is whole-genome *de novo* assembly, although it is less sensitive and computationally very expensive [37].

We recently developed Novel-X [25], a tool for calling novel sequence insertions from SLR sequencing data that shows large improvement over short-read methods. Novel sequences are DNA sequences present in a sequenced sample but missing in the reference genome. Novel-X utilizes barcode information through an assembly of barcode lists originating from the same region. Contrary to the existing short-read methods [13, 18, 17], results were highly concordant with SMRT-SV insertion calls from PacBio long-read data. Novel-X was designed only for novel insertion calling and is not able to call deletion and a wide range of non-novel insertions (e.g. repeat elements).

The development of new algorithms that integrate different types of sequencing data is still in its early stages. In particular, only a few algorithms effectively combine synthetic long-read (SLR) and long-read sequencing data [2, 1]. Here we introduce Blackbird, a novel algorithm for general SV calling that uses a combination of SLR and low-coverage longread sequencing data (hybrid mode). Blackbird provides an SLR-only mode as well. Blackbird is based on the principle of dividing the target genome into small segments, assembling each segment separately using barcode information encoded in the aligned SLRs, and identifying SVs within each segment.

Our novel algorithmic contribution in this paper is two-fold. First, we use interconnecting barcodes assembly in SRL data to solve the general and more challenging problem of SV calling in the 50-10000 bp range. In addition, we implement an assembly graph-based framework to integrate low-coverage long-reads with SLR data to further improve SV detection.

## 2 Methods

The main steps implemented in Blackbird are presented in Figure 1. Below, we describe each step of our method in more detail.

**Figure 1.**
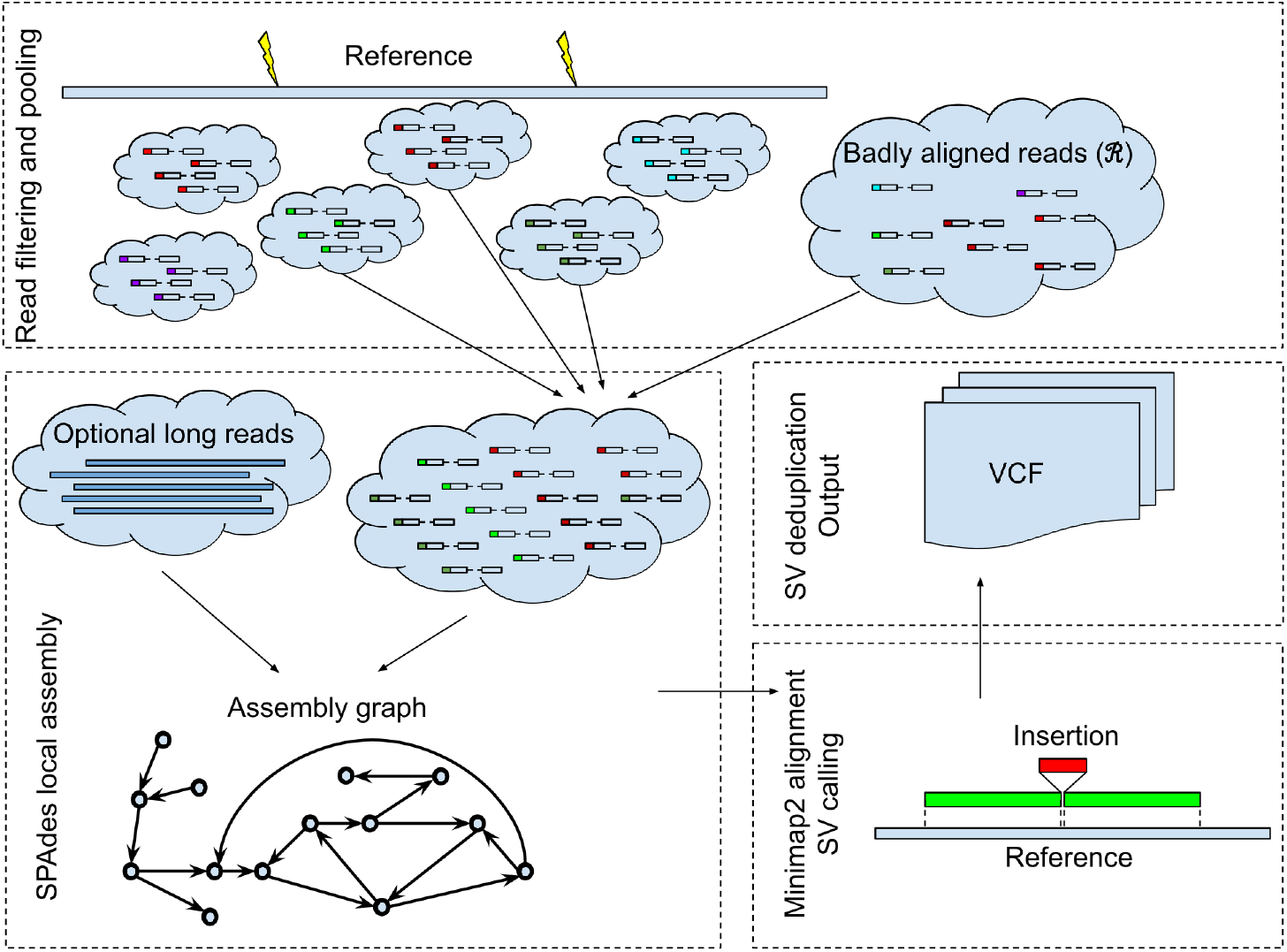
Blackbird outline. The reference genome is split into segments. Inside each segment, we select a set of barcodes and collect reads with these barcodes both aligned into the segment and unaligned. Different barcodes are shown by different colors attached to reads. Then we perform a local assembly for the segment and add long-reads aligned to the segment if they are available. The assembled contigs are mapped back to the reference genome segment using Minimap2, and variants are called from the alignments’ CIGAR strings. At the final step, calls are sorted, deduplicated, and stored in the VCF format.

Blackbird is implemented as a module within the SPAdes software package, leveraging SPAdes utility functions for local assembly. Its primary innovation is a novel approach to efficient processing and filtering, along with the strategic use of barcode information to effectively combine synthetic long-read (SLR) data with long reads.

The input for Blackbird consists of a reference genome (e.g., GRCh38) and an indexed, position-sorted BAM file generated by an aligner that preserves barcode information (we utilize the BX tag of the SAM record for this purpose). We recommend using LongRanger [24] for this task. It is essential for users to employ the same reference genome for both read alignment and as input for Blackbird to ensure consistent results.

### 2.1 Filtering Badly Aligned Reads

First, we search the BAM file for *badly aligned reads* to create a badly aligned read set ℛ. ℛ consists of badly aligned read pairs, that are defined as follows: (a) both reads in pair have a Phred quality higher than 20 for all bases, (b) at least one of the reads is not aligned or aligned to unplaced reference contigs or has more than 20% of soft-clipped bases, or (c) one of the reads has a zero value of AM tag meaning that this read can not be clustered with other reads with the same barcode. Condition (a) is important because we do not want to keep reads of bad quality, that were not ideally aligned because of multiple sequencing errors. Conditions (b) and (c) usually arise when reads that were sequenced around breakpoints position, repetitive regions, and from genome segments missing from the reference genome. We store ℛ in a parallel flat hash map data structure that we refer to as read map ℳ. ℳ internal structure allows to efficiently query a subset of reads from ℛ that have a barcode from a given barcode list ℬ. We refer to a subset of such reads as ℳ(ℬ).

### 2.2 Local Assembly

Then we define a set of segments 𝒮 over the reference genome. By default segments are 50 kbp long and consecutive segments overlap by 10 kbp. Without overlap, events spanning across the borders between segments may not be detected. Conversely, events shorter than 10 kbp will be contained within a single segment and can be detected. As some events may be detected in multiple consecutive segments, we apply a deduplication procedure in later stages to remove redundant calls.

The choice of segment overlap and width parameters is crucial, as it influences the trade-off between assembly complexity and the size of deletions that Blackbird can detect. Our default parameters are designed to minimize the number of repeats in each segment. The length of segment overlap establishes a lower bound on the size of deletions that can be identified; if a deleted sequence completely encompasses the segment overlap, it will not be detected by our algorithm. However, since the majority of structural variants (SVs) are shorter than 10 kbp [6], including notable examples like Alu and L1 insertions and deletions, a segment overlap length of 10 kbp is a reasonable default choice. In this way, we are handling the most challenging classes of insertions and deletions. These parameters can be, however, modified by the user before running the tool if preferred.

At this step, our goal is to assemble one or more contigs that represent the segment, incorporating potential structural variants (SVs) for each segment. During the assembly process, we aim to utilize reads sequenced specifically from this segment while minimizing the inclusion of reads from other segments that may have mapped to it spuriously. For each segment *s* we define a barcode list ℬ(*s*). We want to add barcode *b* to ℬ(*s*) only if a long fragment was taken from *s* and then SLRs were sequenced from the fragment with barcode *b*. Typically, for each correct barcode *b* we can find multiple reads that align in different positions across *s*. Contrary, for incorrect barcodes we expect zero or only a few reads that are aligned locally only in some repetitive regions close to each other. In order to distinguish correct and incorrect barcodes, we use the number of reads with barcode *b* aligned to the segment, and the distance between the leftmost and rightmost reads aligned to the segment. In our experiments, if a barcode has at least 3 read pairs aligned to segment *s* and the distance between the leftmost and rightmost reads is greater than 5*kbp*, we include it in ℬ(*s*), otherwise, it is discarded.

We gather all reads with barcodes in ℬ(*s*) aligned to segment *s* by scanning the position-sorted input BAM-file and storing them as paired reads whenever possible or single reads. Then we add reads from ℳ(ℬ(*s*)). Next, the gathered reads are assembled using our modified version of SPAdes assembler ([28]). We adapted SPAdes to a very specific data type we want to assemble (see Figure 2).

**Figure 2.**
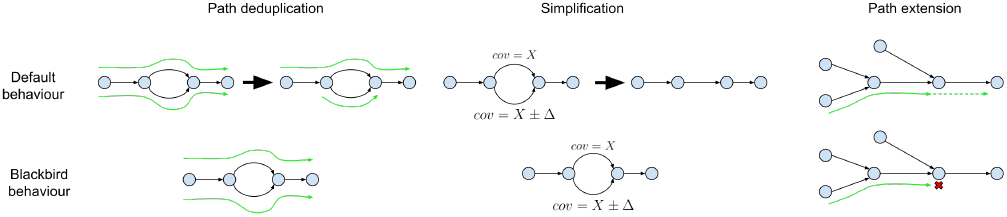
SPAdes adaptation to human segmented data. Left: SPAdes’ path deduplication procedure is relaxed, so if a pair of paths have small differences, both paths are kept for SV calling. **Middle**: SPAdes simplification procedures are relaxed to keep heterozygous SVs. **Right**: SPAdes PathExtend procedure is changed to account for sudden coverage drops, that can form tips in the assembly graph and lead to erroneous deletion calls.

SPAdes was designed for the isolate bacterial data that have completely different properties. First, the human genome is diploid, which creates multiple structures in the assembly graph, that look like bulges in a simple case of heterozygous SNVs. Ideally, for human data, we want to phase these variants, and output two contigs that traverse them. For bacterial data, we do not want to keep these variants because they often arise from very close strains and give minimal information. Such variants, if presented at a low rate, fragment assembly and increase duplication ratio, which is harmful to downstream analysis. Originally in SPAdes, two procedures were implemented to make duplication ratio close to one. First, simplification procedures aggressively remove tips and bulges even if their coverage is high. In the case of bulges, an edge with lower coverage is removed. Second, at the last SPAdes step path deduplication procedure is performed. During this procedure, if two paths are almost identical (e.g. threading through a different side of a bulge), they are deduplicated. A path of a higher coverage is kept intact, and for the second path, all overlaps with the higher-covered path are removed. For SV calling problem, high duplication ratio does not affect precision or recall because paths that would be deduplicated are very similar to other paths, and the same structural variant would have been called from them. Such variants can be deduplicated in later stages. Therefore in order to adapt SPAdes for the human data, we disabled path deduplication procedures, and relaxed simplification procedures to keep variation aroused due to the diploid nature of the human genome (see Figure S1).

In [29], the authors present examples of misassemblies in assembly graphs that appear error-free. These misassemblies are often attributed to imperfect genome coverage resulting from sequencing biases. Even a local drop in coverage can lead to a misassembly event, particularly in repetitive regions. Unfortunately, SLR data frequently exhibit sequencing biases that create coverage gaps across the genome (see Figure S2).

Mapping-based methods can typically distinguish coverage drops from actual deletions, as there are no split-read or discordant paired-end read signatures associated with the coverage drop. However, for assembly-based methods like SVABa or Blackbird, coverage drops in repetitive regions may result in false positive deletion calls.

To mitigate this issue, we implemented a new topology rule for contig extension (Figure 2). In scenarios with perfect coverage of the target genome, it is safe to extend a path using an edge without alternatives. However, coverage drops can create long tips, which we do not expect to encounter in human sequencing data. We modified this rule to check for the presence of long tips in the vicinity and refrain from extending the contig in such cases.

In hybrid mode, long-reads are also used during the local assembly step. We add those long-reads that are aligned to the segment to the assembly. Path extension procedure for hybrid assembly was not altered, but we added long-read sequences to the graph construction procedure. Another improvement was to use long-reads to close the gaps not only between tips but also between arbitrary parts of the graph. In this gapclosing step, we limit the number of long-reads to 100 reads per segment to reduce the computational cost (see Figure S3).

### 2.3 Identification of Structural Variants

Assembled contigs are aligned to the reference genome using minimap2 ([21]). Insertions and deletions are called using the CIGAR string of minimap2 alignments. Due to the diploid nature of the genome, some events might be called multiple times. We implemented a deduplication procedure that removes events that have the same breakpoint location and length as some other event. Resulting insertions and deletions are sorted and stored in a VCF file.

### 2.4 Software availability

Blackbird is freely available at https://github.com/1dayac/Blackbird.

## 3 Results

Benchmarking SV detection tools is quite challenging due to the large number of available tools and the difficulty of selecting a subset of relevant ones. For comparison, we picked Manta [7] since according to [19], a recent comprehensive evaluation of SV detection tools, this is the best short-read method overall on the real data. As Blackbird is an assembly-based method, with a focus on smaller and medium-sized events, we also selected SvABA [35] as a representative comparison. LEVIATHAN [26] is an SLR SV detection tool that uses barcode information, while LongRanger is an SLR aligner that calls deletions during the read alignment step. LongRanger is not able to call insertions and it was only assessed for deletions in our evaluations.

To the best of our knowledge, there are no existing SV callers that are able to work in this hybrid setting. We decided to compare Blackbird in hybrid mode against the state-of-the-art long-read SV callers - Sniffles2 [30] and PBSV.

We compared Blackbird, long-read, and short-read tools performance on the simulated dataset, and the HG002 dataset that has a well-established benchmark callset of insertions and deletions [39].

We tested Blackbird in SLR mode against Manta, SvABA, and LEVIATHAN, and in hybrid mode against Sniffles and PBSV. Specifications of tested tools can be found in “Supplementary text A: Tool specifications”. For each tool, we compare calls with insertion/deletion sizes between 50-10000 bp. For SV matching and comparison we used Truvari [10], which allows us to compare a callset obtained by one of the tools to the reference callset. First, we divide target and reference VCF files into separate deletions and insertions/duplication VCF files. These files are compared against each other using Truvari. We exclude events shorter than 50 bp because all assembly-based methods have low precision for short events since genome assembly is prone to mismatches and small insertions and deletions. Precision and recall are the metrics we use to evaluate methods. The exact command we used to run Truvari can be found in “Supplementary text B: Running Truvari”.

### 3.1 The Simulated dataset

### 3.2 Hybrid mode

In order to evaluate the performance of Blackbird within a fully controlled environment, we first created an SLR simulated dataset. We added insertions and deletions from [16] to chromosome 1 of the GRCh38 reference genome. Such an approach allows us to test performance on various insertion and deletion repeat families, such as L1, *Alu*, and STR. We inserted 1,437 events with a total length of 799 kbp. Additional information about these events can be found in “Supplementary text C: Simulated SV events”. We then simulated a 37× SLR dataset using LRSIM [23] with default read simulation parameters. We aligned reads to chromosome 1 of the GRCh38 genome using LongRanger. Additionally, we simulated PacBio HiFi long reads with 9000-12000 read lengths and 0.01-0.001 error rate per read, at coverage levels of 5, 10, 15, 20, 25, and 30, using bbmap’s randomreads module [3]. We tested Blackbird in Hybrid mode against state-of-the-art long-read SV callers PBSV and Sniffles. We compare the results of the methods above with the list of ground-truth SVs. We count a predicted SV as correct if there exists a corresponding SV in the ground truth set of the same type and their position on the reference is not further than 100 bp away from each other. Blackbird was able to achieve 85-87% recall for deletions, 75-77% recall for insertions, and precision in the 90-93% range for all datasets (Figure 3). Using these results we can argue, that SV callers have saturation property, such that at some point additional long-read coverage adds only incremental or no improvement to the results. Blackbird’s results in hybrid mode are superior than the results in SLR mode even with 5× long-read coverage, but results do not improve with increased long-read coverage. Sniffles performance saturates at 10× long-read coverage and PBSV does not saturate at lower than 10× rate but saturates at 15× rate. It can be seen that Blackbird in hybrid mode with 5× long read coverage is superior to PBSV and Sniffles with 5× coverage, and comparable (slightly better for insertions and slightly worse for deletions) with Sniffles, PBSV with 15× coverage.

**Figure 3.**
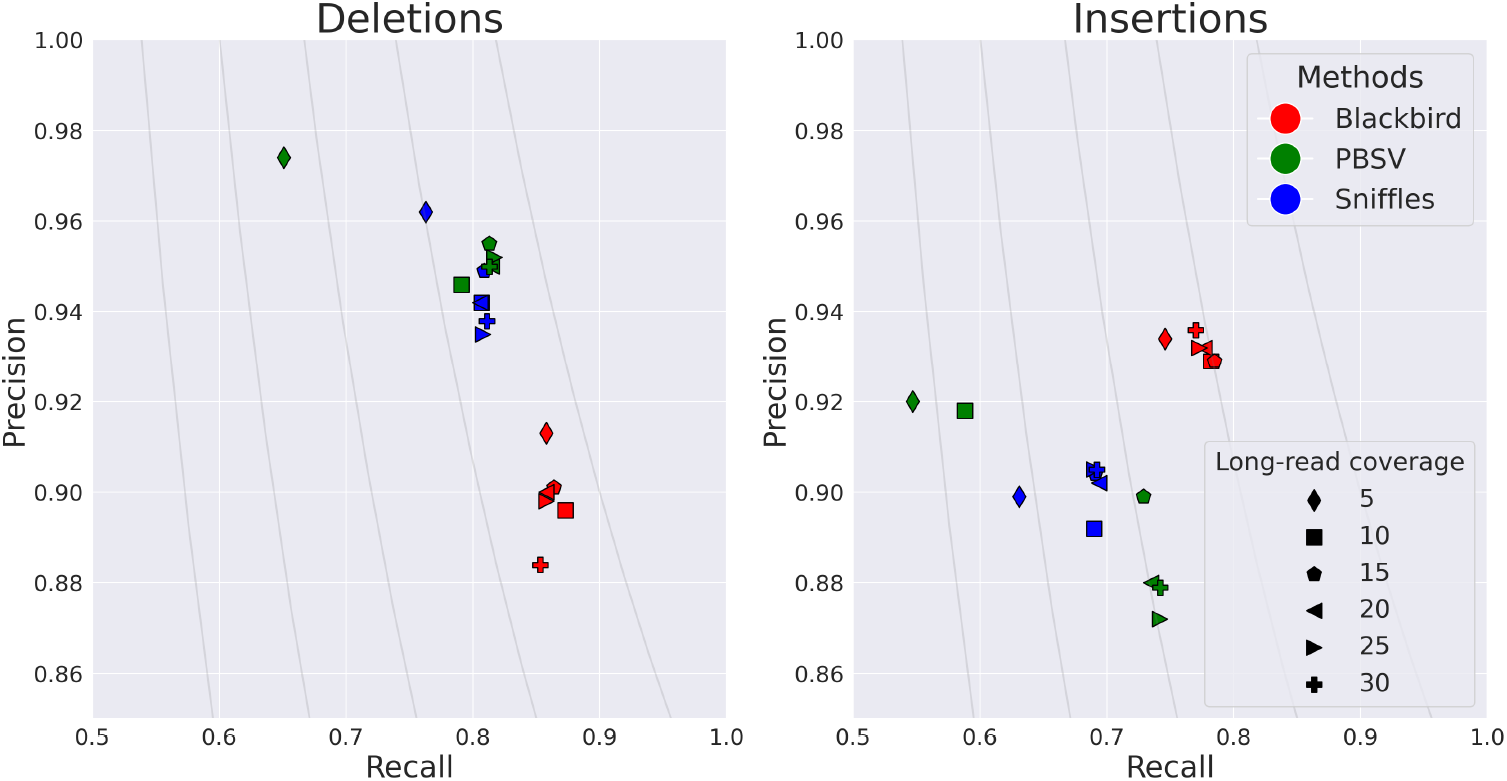
Results for the hybrid simulated dataset. Blackbird is able to achieve the highest precision and recall for insertions, and better recall for deletions. While Blackbird has lower precision for deletions than competitors, it is able to achieve better F1-scores. Grey curves are F1 curves.

These findings lead to the conclusion that Blackbird can achieve the same SV calling quality with lower long-read coverage. Thus, Blackbird can be a cost-effective alternative or can be used when high coverage with long reads is not readily available.

### 3.3 SLR mode

We evaluated the performance of Blackbird, LongRanger, Manta, SvABA, and LEVIATHAN on the same simulated SLR-only dataset. We show the precision and recall of each method in Figure 4.

**Figure 4.**
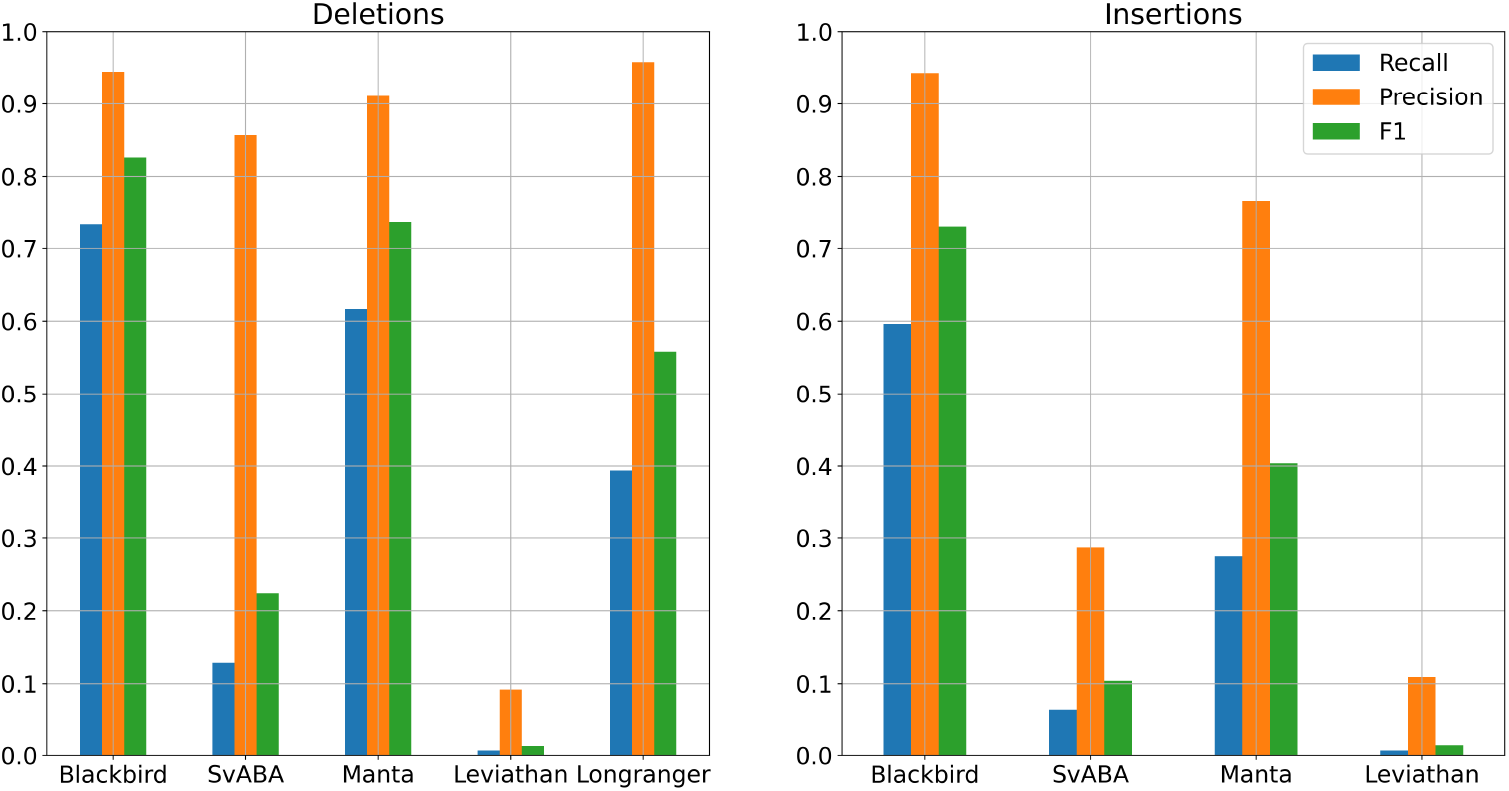
Results for the SLR simulated dataset. Blackbird is able to achieve the highest recall and F1-score both for insertions and deletions. These results show that Blackbird is able to call insertions with much higher recall that other short-read SV calling tools.

On the simulated dataset, Blackbird achieved the best overall result with the highest sensitivity and F1 score for both insertions (58% and 0.71 respectively) and deletions (72% and 0.81 respectively). Additionally, Blackbird achieved the same precision for insertions (97%) as Manta and SvABA, and managed to obtain precision for deletions at 93% level, slightly behind LongRanger (95%) and SvABA (96%). Therefore, Blackbird achieves the best average results on the simulated dataset. Interestingly, the performance of all short-read/SLR tools is limited by a majority (54%) of STR SVs in the simulated data. Blackbird is also able to call the most of *Alu* repeats and half of L1 repeats.

### 3.4 The HG002 dataset

The HG002 genome has been extensively characterized by the Genome In A Bottle (GIAB) consortium with a curated insertion and deletion benchmark callset [39] for the GRCh37 reference genome. We used UST TELL-Seq data with 42× coverage and 169 bp read length with a 20% addition of shorter reads and stLFR data with 35× coverage and 100 bp read length as benchmark SLR datasets. These two technologies are among the most popular SLR technologies publicly available. We aligned SLR reads to the reference genome using LongRanger that was modified to accept TELL-Seq and stLFR barcodes. For PacBio HiFi data, we used a dataset from the PacBio website with an average read length of around 16500 bp for 10× and 5× coverage, and T2T PacBio HiFi data with an average read length of around 12500 bp for 34× and 20× coverage. In order to get 5× and 20× datasets, we downsampled higher-covered PacBio datasets. For SLR mode comparison we used the stLFR dataset mentioned above. The results are presented in Figure 5 and Figure 6. In the SLR mode comparison, Blackbird shows comparable recall with Manta on deletions, but has the highest recall for insertions, while keeping reasonable precision. This result shows the advantage of SLR technologies over standard short-read sequencing demonstrated by our algorithm. Blackbird was able to call long insertions exceeding 1000 bp in size. In hybrid mode comparison, PBSV and Sniffles have a stable precision rate, and recall increases with increased coverage. However, it is clear that on the HG002 dataset results saturate only with 10-20× long-read coverage. When long-read coverage drops to 5× rate, precision drops by 10-15% for both insertions and deletions compared to 10× coverage. On the other hand, when TELL-Seq data is used, Blackbird’s recall drops only by 4% for deletions and 2% for insertions. At 10× coverage, Blackbird also performs better in terms of recall compared to Sniffles and PBSV. Blackbird with stLFR data performs worse than Blackbird with TELL-Seq data when the long-read coverage is within 5-10× range but achieves the best recall compared to all other methods on higher long-read coverage. These results show that Blackbird, especially with TELL-Seq data, is able to achieve saturation earlier than Sniffles and PBSV. They also suggest that for the SV detection problem, there is no need to go deeper than 20×, as the improvements are only incremental at higher coverage.

**Figure 5.**
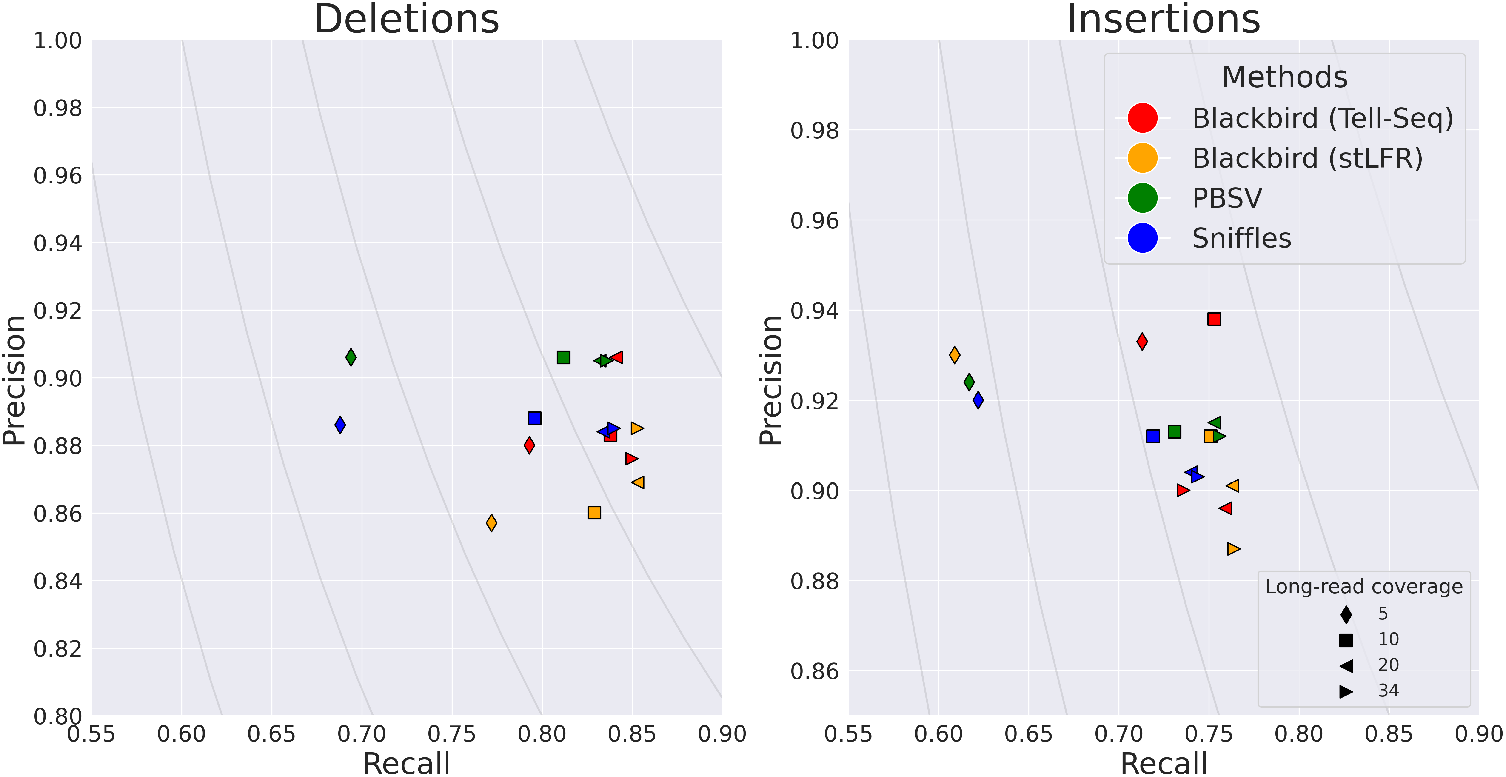
Results for the HG002 hybrid dataset. Left: Deletions. Blackbird results saturate with lower long-read coverage. E.g. Blackbird on Tell-Seq data and stLFR data with 5× long read coverage appear close to the runs with the higher coverage on the plot. PBSV and Sniffles have a 12-15% drop in recall when long-read coverage is equal to 5×. Right: Insertions. Blackbird on Tell-Seq data performs similarly to the deletions run. However, Blackbird on stLFR data with 5× long-read coverage performs similarly to Sniffles and PBSV. Also, Blackbird is able to achieve the highest recall for insertions and deletions when the coverage is high. Grey curves are F1 curves.

**Figure 6.**
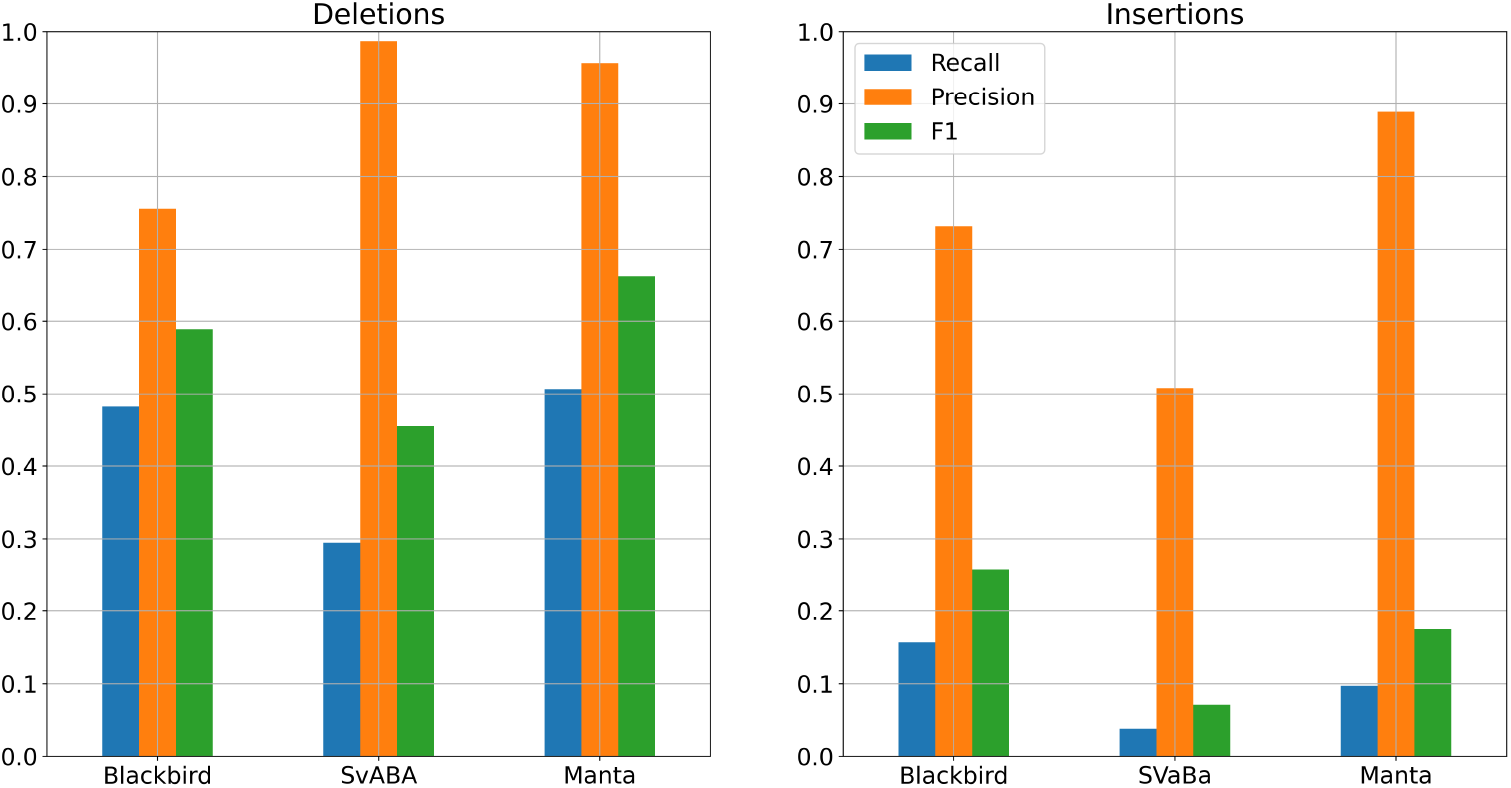
Results for the SLR HG002 dataset. Left: Deletions. Manta achieved better recall and precision for deletions, though Blackbird recall is only 2% lower. Right: Insertions. Blackbird is able to double the recall for the insertions on the HG002 dataset compared to the Manta. Some of the insertions found by Blackbird were longer than 1 kbp.

We explored what kind of sequences are called with Blackbird in hybrid mode with TELL-Seq and long reads at 5× coverage but are not called by Sniffles and PBSV at 5× long-read coverage (see Figure S4). In order to do that, we extracted SV sequences from the GIAB callset, ran them through RepeatMasker, and counted the amount of called sequences for each reported repeat type. Results show that there is no designated repeat type that benefited exclusively. Additional Blackbird calls are scattered across different repeat types and non-repetitive sequences uniformly, excluding simple repeats, where the number of calls is almost similar. Information about how zygosity and long-read coverage impact recall can be found in “Supplementary text D: Zygosity and Long-Read Coverage in SV Calling”.

### 3.5 Dataset of challenging medically relevant genes

Another source of curated SV calls is the dataset of challenging medically relevant autosomal genes [34] for the HG002 genome. This dataset contains around 400 SVs, which are reported to be more challenging on average than the variants from the GIAB dataset. We performed additional benchmarking using this dataset. The results can be found in “Supplementary text E: Dataset of medically relevant genes” which further confirm the utility of our method.

### 3.6 Running time analysis

We conducted experiments to assess the running time and memory usage of Blackbird on Intel Xeon Gold 6230 2.1-3.9GHz processors with 32 cores, using the HG002 dataset. For our SLR dataset, we used the 42× stLFR dataset mentioned earlier, and for our long-read dataset, we used the 34× PacBio HiFi dataset.

Blackbird, when operating in SLR mode, was expectedly slower than the standard short-read methods SvABA or Manta. Specifically, on the HG002 dataset, Blackbird required 6 hours and 12 Gb of memory, while SvABA finished in 65 minutes and 11.5 Gb and Manta was able to complete the task in just 15 minutes and 6.5 Gb.

When running in hybrid mode, Blackbird took 11 hours to finish and utilized 117 Gb of memory. In comparison, Sniffles2 required only 3 minutes and 0.6 Gb, while PBSV needed 1 hour 19 minutes and 27G. Despite these performance differences, we believe that Blackbird’s increased sensitivity with low-coverage long reads makes it a viable alternative for SV calling.

## 4 CONCLUSION

We introduced Blackbird, an efficient and sensitive approach for SV calling. While the current implementation of Blackbird is limited to insertions and deletions, we plan to implement modules for inversions in the future. While Blackbird is not capable of working with larger events (e.g. translocations), there are available tools that can call that events effectively. We also plan to improve the performance of our algorithm on diploid data, such as distinguishing between homozygous and heterozygous variants and adding variant phasing. At the moment Blackbird operates on a contig scale, which makes haplotyping and homozygous/heterozygous classification challenging. However, our integration with SPAdes genome assembler makes it possible to use an assembly graph structure to better classify heterozygous variants.

Blackbird assembles each genomic segment, even regions where readpair or split-read signal cannot be found. On the other hand, potentially Blackbird can be transformed into a reference-assisted SLR assembly tool. There are no existing algorithms that solve this problem, and Blackbird’s performance is already better than existing *de novo* assembly tools such as Supernova. Future work can also include running time optimization. A major bottleneck is caused by the need to constantly search for read mates in a large indexed BAM file. BAM index provides semi-random access by position, its running time is highly dependent on the local coverage around the mate’s position. For some segments, the naive solution, that requires a separate query for each mate can be too slow.

## 5 Data access

- **Tell-Seq Data** is available under SRX7264481 accession in SRA.
- **stLFR Data:** Link to stLFR Data
- **PacBio Hi-Fi Data:** Link to PacBio Hi-Fi Data
- **ONT Data** is available under PRJNA200694 BioProject

## Acknowledgement

This work was supported by the National Institute of General Medical Sciences (NIGMS) of the National Institutes of Health (NIH), Maximizing Investigators’ Research Award (MIRA) R35GM138152 to I.H. The content is solely the responsibility of the authors and does not necessarily represent the official views of the National Institutes of Health.

