## Supplemental Table for "Blackbird: structural variant detection using synthetic and low-coverage long-reads"

#### Supplementary text A: Tool specifications

- Blackbird - build from 50cf64605f776ad2ca976f385e65d4210f864ba0 commit (September 9, 2022)
- Manta - v1.6.0 (`https://github.com/Illumina/manta/releases/tag/v1.6.0`)
- SVaBa - v1.1.0 (`https://github.com/walaj/svaba/releases/tag/1.1.0`)
- LEVIATHAN - v1.0.1 (`https://github.com/morispi/LEVIATHAN/releases/tag/v1.0.1`)
- Long Ranger - v2.1
- Sniffles - v2.0
- PBSV - 2.6.2

#### Supplementary text B: Running Truvari

For simulated experiment:

```
truvari bench -c <input_vcf> -b <ground_truth_vcf> -o <output_folder>
    --pctsim=0
```

For HG002 experiment:

```
truvari bench -b <input_vcf> -c <ground_truth_vcf> -o <output_folder>
    --includebed HG002_SVs_Tier1_v0.6.FINAL.bed --pctsim 0 --sizemax=10000
    --sizemin=50 --sizefilt=40 --refdist=500 --pctsize=0.5 --pctov=0.5
    --typeignore
```

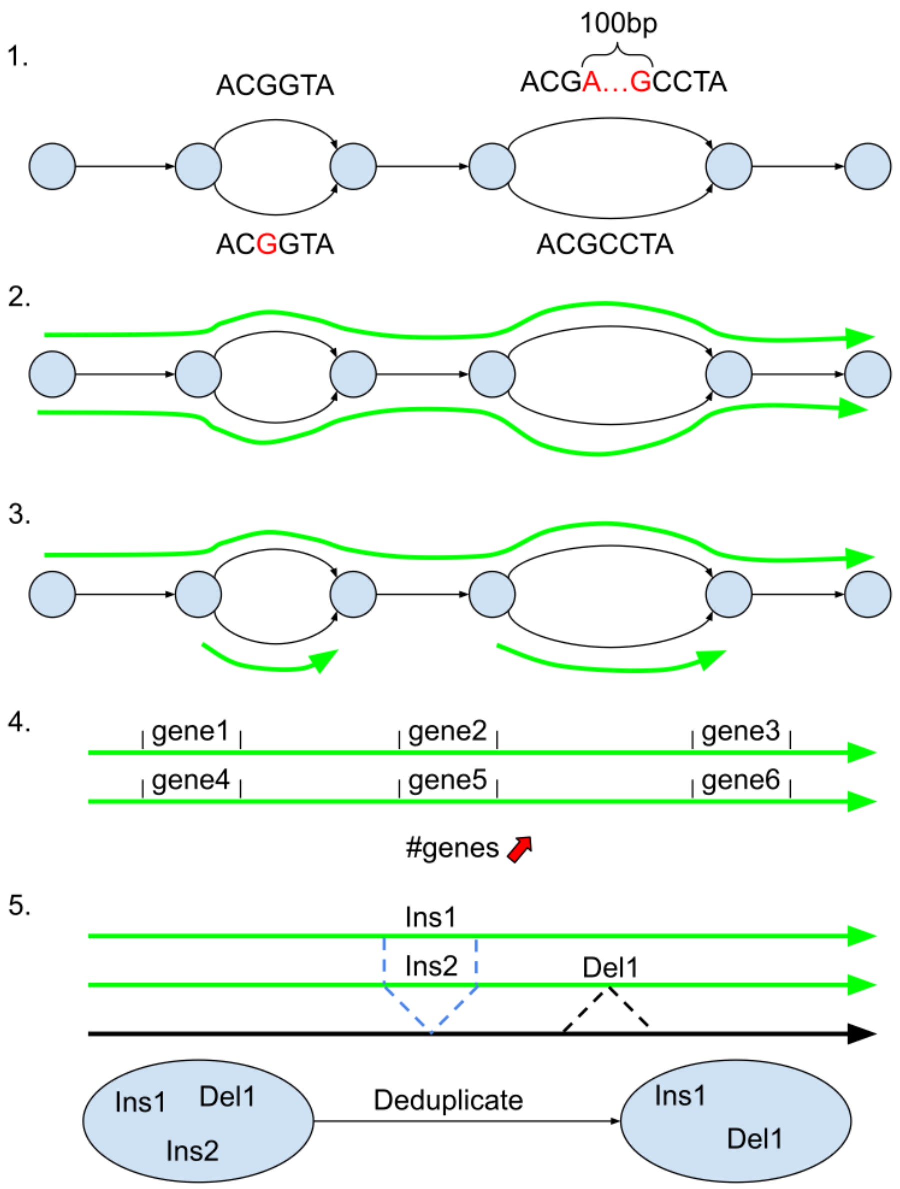


Figure S1: **SV calling and genome assembly duplication ratio.** 1. An example of an assembly graph with an SNP on the left, and 100bp insertion on the right. 2. Blackbird's version of SPAdes will likely to generate two long contigs from this assembly graph. 3. SPAdes in default mode will generate one long, and two short contigs. 4. The reason for default SPAdes behavior is that most downstream applications (e.g. gene prediction) require duplication rate close to 1. In this example, gene count will be 2× inflated because some kind of variation is presented in the sample. 5. For SV detection inflated duplication ratio is not crucial, because it is easy to deduplicate SV calls.

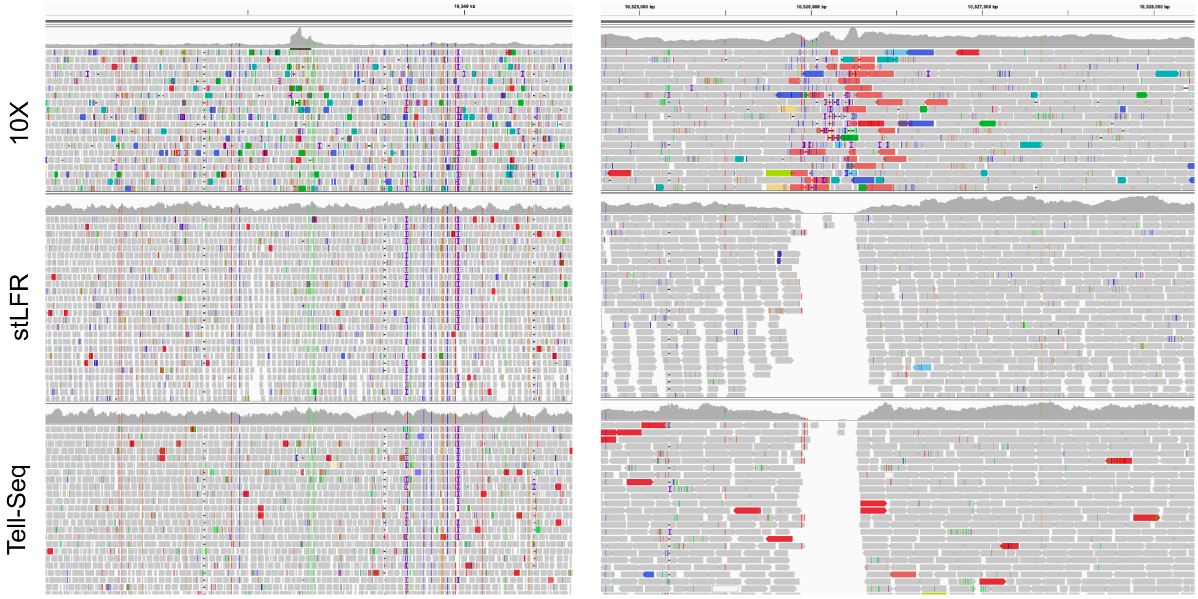


Figure S2: **Properties of different SLR technologies.** Three popular SLR technologies have different data properties. 10X Genomics SLR data is known to have short overamplified regions. E.g. on the left figure 10X data has coverage up to 750 and can be up to tens of thousands in other regions. Blackbird's filtering step allows for reducing coverage inside these regions. stLFR and Tell-Seq often have coverage gaps in repetitive regions (right figure). While Blackbird in SLR mode can't fill these gaps even using barcode information, long reads in hybrid mode allow to close them effectively.

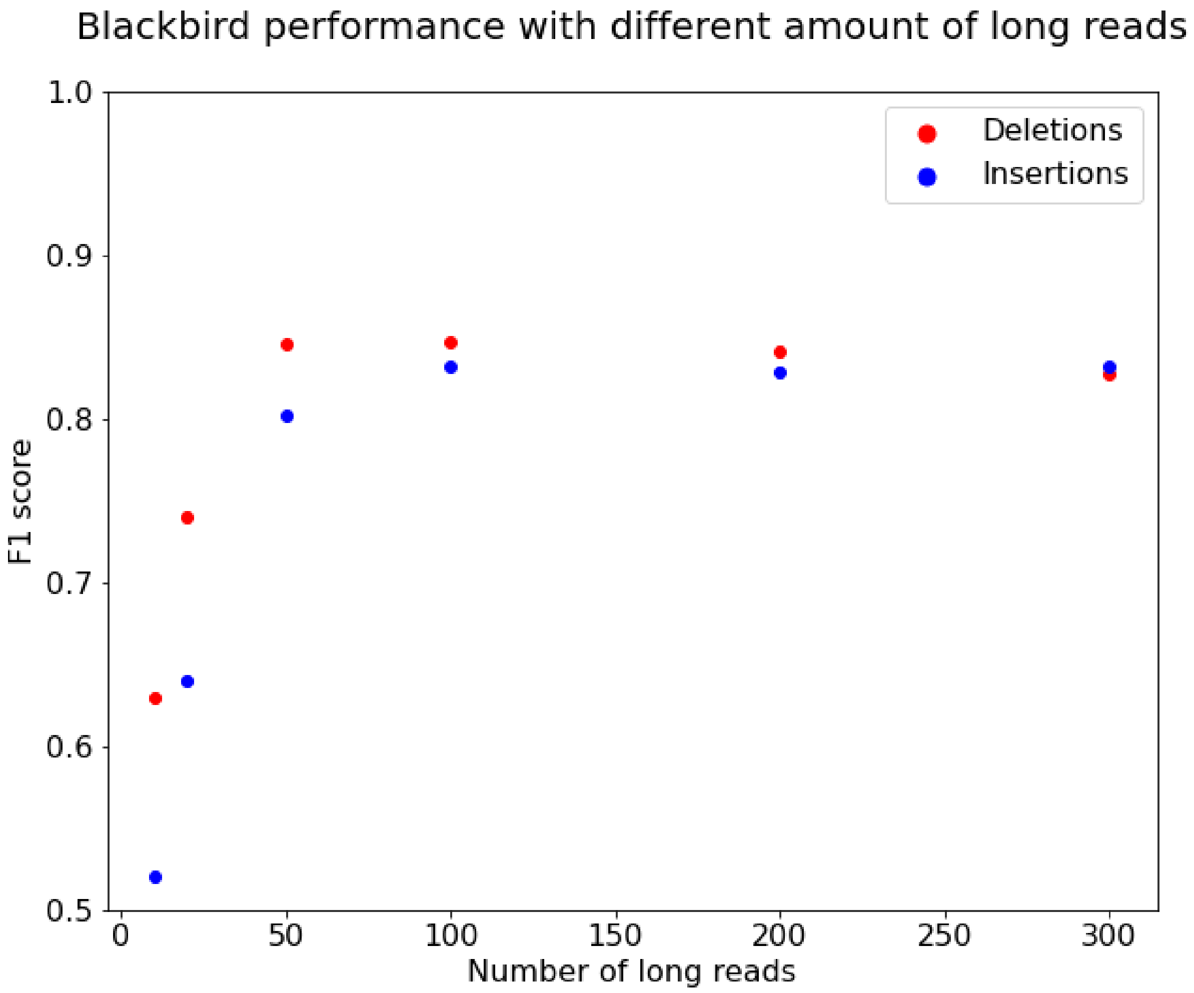


Figure S3: **Blackbird SV calling performance on HG002 dataset depending on the number of long reads.** We can see that F1 scores stop increasing for insertions when we have 100 long reads, while deletions require only 50 long reads to saturate. Therefore we chose 100 as a cutoff for the number of long reads per local assembly, since long read alignment in SPAdes is computationally expensive.

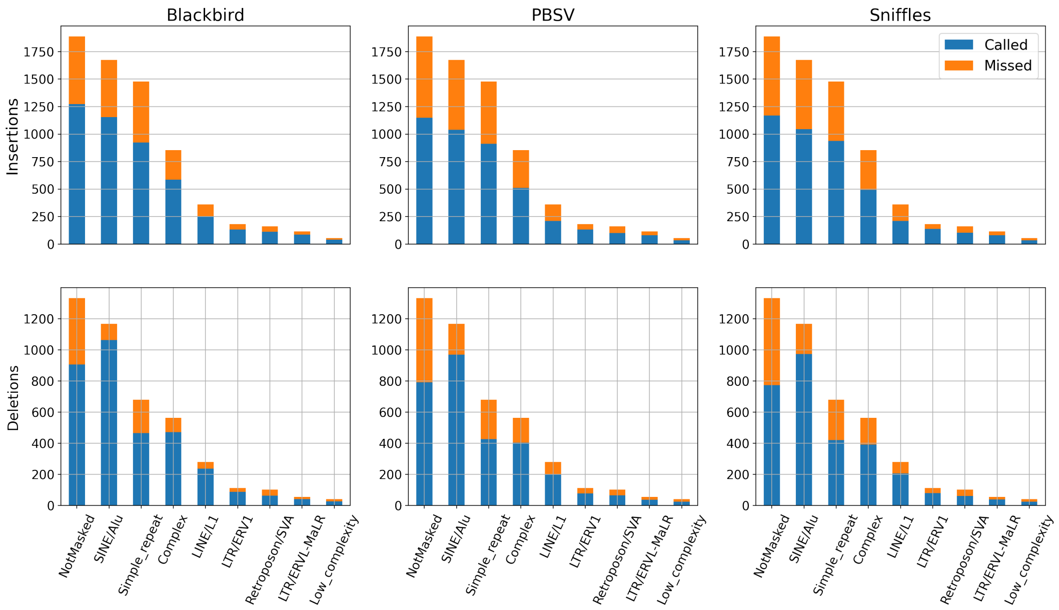


Figure S4: **Repeat type breakdown for the HG002 dataset.** This plot shows the repeat type breakdown and the number of events called at 5× coverage for different tools. These plots do not show an obvious trend of increased recall vs repeat type. Blackbird uniformly calls more events than the competitors, regardless of the predicted repeat type.

### Supplementary text C: Simulated SV events

To test the performance of different SV calling tools, we created an artificial chromosome with insertions and deletions from [1] using a custom script. We selected the largest human chromosome (chromosome 1) to allow for quicker result turnaround, but kept the number of simulated events high to ensure adequate coverage of various repeat families, such as L1, *Alu*, and STR.

In total, we inserted 1,437 events with a total length of 799 kbp. The distribution of event types, lengths, and positions can be found in Figure SS1.

### Supplementary text D: Zygosity and Long-Read Coverage in SV Calling

We evaluated the performance of SV calling tools by comparing the impact of zygosity and long-read coverage on recall. We divided the HG002 Tier 1 callset into four subsets: heterozygous deletions, heterozygous insertions, homozygous deletions, and homozygous insertions, and estimated recall for each event type with varying long-read coverage. Note that this procedure omits some SV calls, because not every event in the Tier1 callset has an unambiguous homozygous or heterozygous genotype.

We ran Truvari and calculated how many events were called by each tool on different coverage for each subset, and plotted results in Figure SS2. This Figure shows that Blackbird, Sniffles, and PBSV have different patterns of SV calling across different SV types and coverages. So Blackbird shows the best results at 5× coverage across all SV types. For high long-read coverage, Sniffles shows better results for heterozygous deletions, while PBSV achieves better results for other types of events. In conclusion, the results of our experiment show that each SV calling tool has its own unique strengths and performs best in different scenarios, though the results of each tool are relatively close to each other and highly concordant.

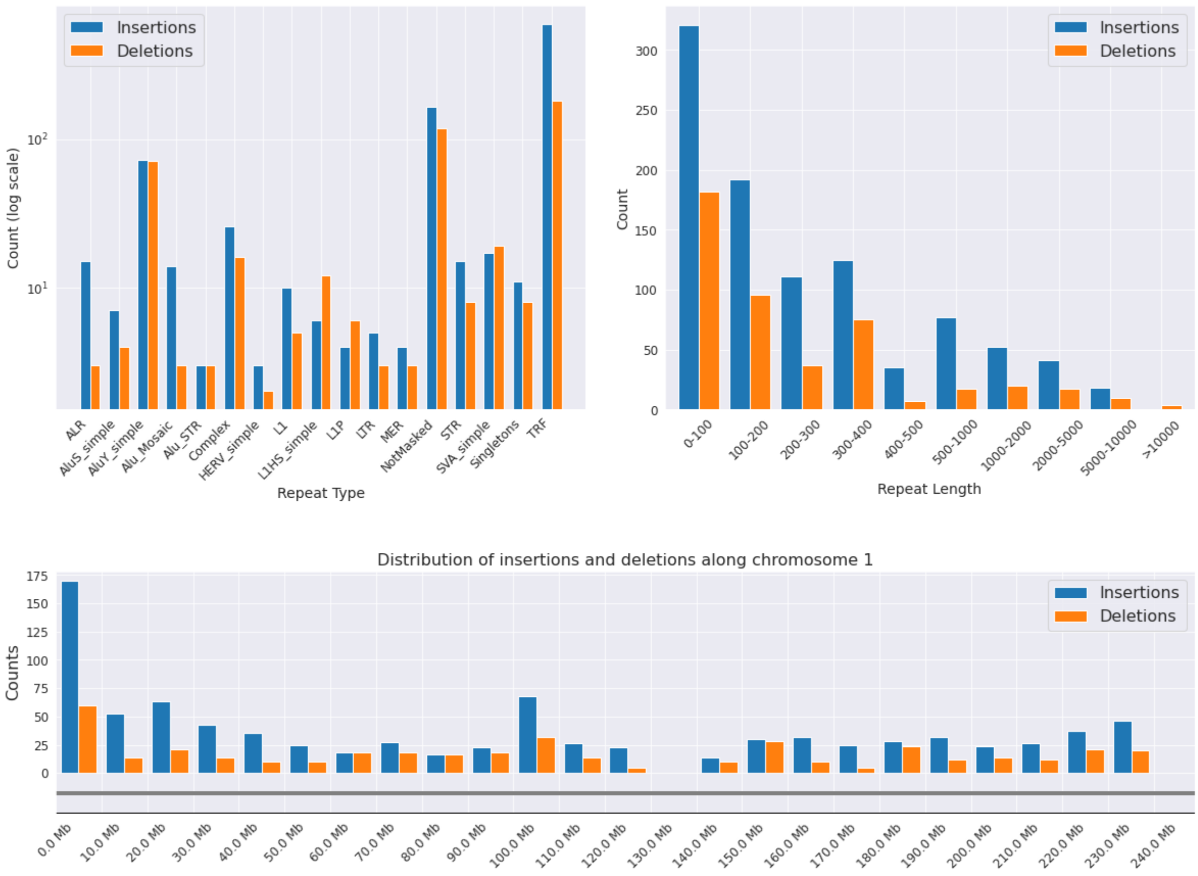


Figure SS1: **Properties of SV events in the simulated dataset. Top left:** The frequency of repeat types in a set of simulated insertion and deletion variants. Tandem repeat elements are the most common repeat type, followed by non-repetitive sequences and simple ALU repeats. **Top right:** The distribution of lengths in the simulated insertion and deletion variants. Most insertions and deletions are less than 1000 base pairs in length, with a smaller number of longer variants. **Bottom:** The distribution of simulated insertion and deletion variants across chromosome 1 of the human genome.

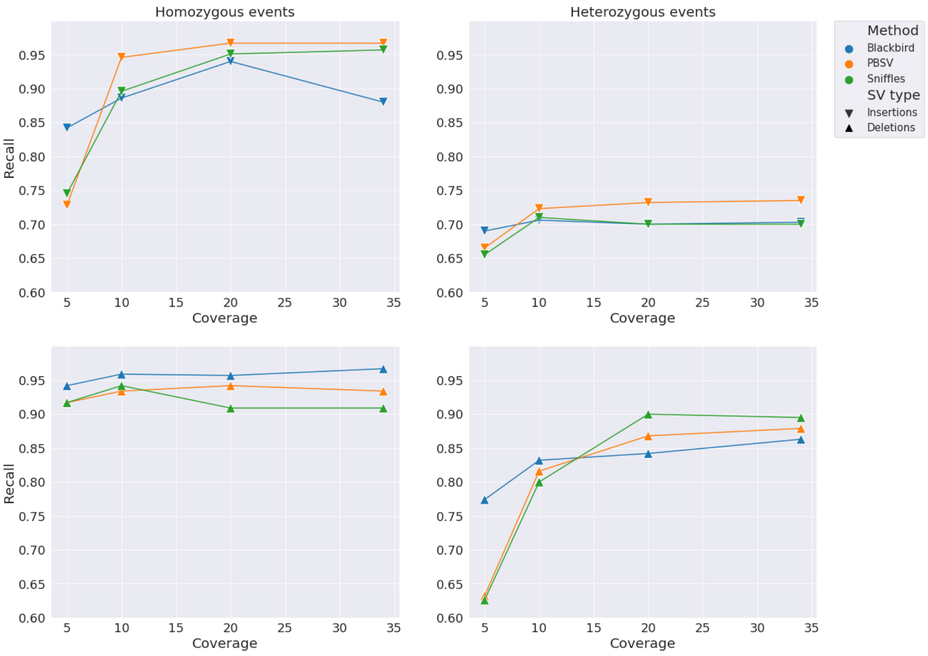


Figure SS2: **Comparison of recall for different methods and SV types in homozygous and heterozygous events.** The plot illustrates that recall is higher for homozygous events than for heterozygous events, and that recall generally increases as coverage increases. Additionally, the method Blackbird is shown to have higher recall at lower coverage levels.

### Supplementary text E: Dataset of medically relevant genes.

The dataset from [2] comprises approximately 200 curated SVs, which are considered to be more complex than their counterparts in the GIAB Tier 1 dataset. We conducted benchmarking of Blackbird in hybrid mode using stLFR SLR data, Sniffles, and PBSV on this dataset using a strategy identical to that employed for the HG002 GIAB callset. Precision and recall values for various coverage levels are depicted in Figure SS3. These results show that for this dataset Blackbird results are slightly worse than PBSV or Sniffles in terms of precision, though recall values are very similar.

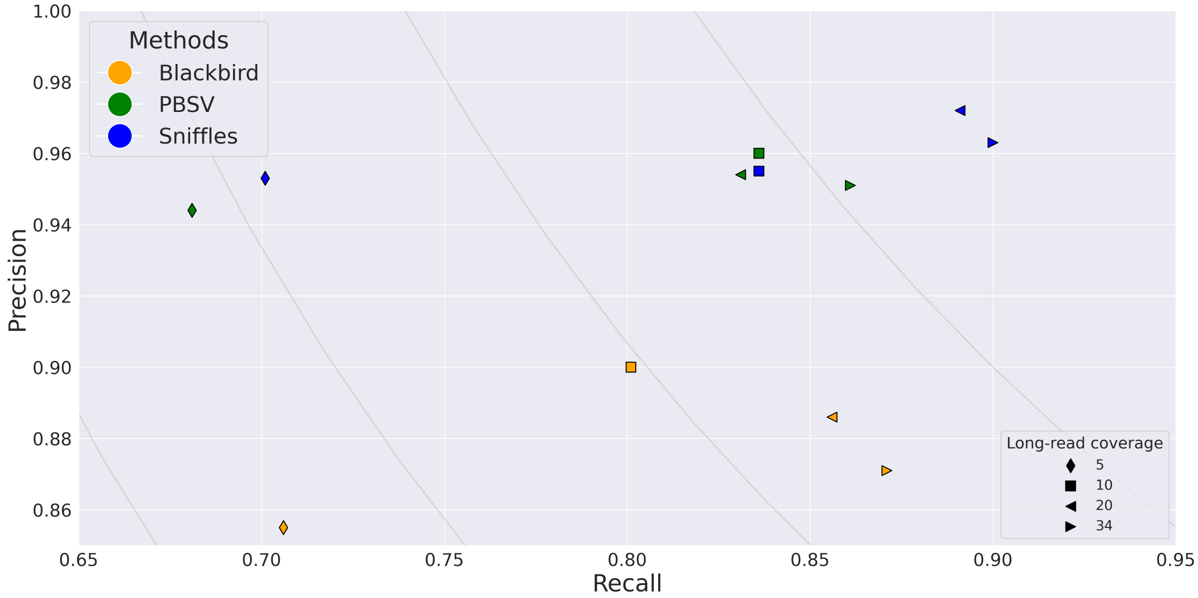


Figure SS3: **Benchmarks on medically relevant genes dataset.**
